# A repository of web-based bioinformatics resources developed in India

**DOI:** 10.1101/2020.01.21.855627

**Authors:** Abhishek Agarwal, Piyush Agrawal, Aditi Sharma, Vinod Kumar, Chirag Mugdal, Anjali Dhall, Gajendra P.S. Raghava

## Abstract

IndiaBioDb (https://webs.iiitd.edu.in/raghava/indiabiodb/) is a manually curated comprehensive repository of bioinformatics resources developed and maintained by Indian researchers. This repository maintains information about 543 freely accessible functional resources that include around 258 biological databases. Each entry provides a complete detail about a resource that includes the name of resources, web link, detail of publication, information about the corresponding author, name of institute, type of resource. A user-friendly searching module has been integrated, which allows users to search our repository on any field. In order to retrieve categorized information, we integrate the browsing facility in this repository. This database can be utilized for extracting the useful information regarding the present scenario of bioinformatics inclusive of all research labs funded by government and private bodies of India. In addition to web interface, we also developed mobile to facilitate the scientific community.

## Introduction

Bioinformatics era beginning is marked by the development of database Protein Information Resource (PIR) by Margaret Dayhoff in the late 1960s [1]. The database comprises of a large number of protein sequences and its analysis for studying molecular evolution. Further, scientists developed numerous algorithms for analyzing the available biological data. Bioinformatics in India marks its start with the discovery of the famous Ramachandran plot that establishes the foundation of modern structural biology/bioinformatics [2]. As a result of these efforts, bioinformatics has grown full-fledged in the Indian study curriculum and research. This effort was further boosted by the establishment of the Biotechnology Information System Network (BTISNet) by the Department of Biotechnology (DBT), Govt. of India in the 1980s. Several centers were set up across the country under BTIS with the goal to carry out research in this area and providing services to the scientific community. These centers are further distributed into various levels such as Centers of excellences (CoEs), Distributed Information Centers (DICs), Distributed Information sub-centers (Sub-DICs), Bioinformatics Infrastructure Facilities (BIFs) [3]. BTIS has provided state of the art of the computational infrastructure to the various research institutes and universities for promoting scientific research as well as assisted several research and development projects by creating and maintaining databases, computational algorithms, and webservers.

High throughput studies in biology and medicine have generated a large amount of data for analysing genomes, transcriptome, epigenome, metabolome, and microbiome [4]. Development in mass spectrometry techniques has generated enormous amount of proteome data, which together constitutes a major portion of biological data. Datasets related to protein-protein interactions, protein-small molecule interactions, protein-nucleic acids have been compiled through years due to advances in techniques like NMR, X-ray, SAXS, Cryo-electron microscopy, etc. Therefore, data-driven research has been one of the important processes in the last few decades to solve biological questions. India is one of the few countries which has acknowledged the increase in the volume of sequence and structural data in the field of biology and medicine. Scientists in India have developed several biological databases and software for data analysis and visualizations to solve biological problems. India is currently gaining huge attention for generating a massive amount of new biological data in the house as well as for developing a number of computational resources. This can be clearly seen at website of Database Commons; India is one of the leading countries and stands at 4^th^ position after the USA, China, and the United Kingdom in the area of developing and maintaining biological databases (http://bigd.big.ac.cn/databasecommons/stat#).

Most of the popular scientific journals published by different publishing groups (e.g., Nature publishing groups, Elsevier, Oxford publishers) are dedicated for bioinformatics resources. This clearly indicate the importance of bioinformatics resources in the field of biological science. In the past, most of the research articles published on bioinformatics in India [5–10], reviewed Indian education system and computational resources developed by Indian researchers. Best of our knowledge, there is no website, portal or database that maintain links to bioinformatics resources developed by Indian researchers. This study complement previous studies and facilitate scientific community in accessing web-based resources developed by Indian bioinformaticians. In this article, we described a web-based platform and mobile app developed to maintain web-based services (prediction methods and databases) developed by Indian community. Similar type of analysis has been performed for in the past for different countries [11–15].

## Material and Methods

### Data Collection and Organization

We manually collected and curated the data from PubMed by searching keywords like “database”, “prediction”, “webserver”, “*in silico* study” and “http/https” in the Title/Abstract section and “India” in Affiliation section up to June 2019. The search resulted in around 750 articles. We read all these articles and select only those articles which provide the link to a webserver. We excluded those entries in which the web servers were not functional or the authors are Indian however servers are maintained outside the India. Finally, we got 541 research articles which describes a functional webserver. Comprehensive information was extracted from these papers that includes “Name of server”, “Description”, “Link”, “Journal”, “Year”, “PMID”, “Author”, “Email”, “City”, “Status”, “Category”, and “Class”.

### Development of Web-based Platform

All information extracted from relevant research papers were curated wisely to create a database. This database was developed using MySQL which is a free relational database management system. In order to launch web service Apache was installed and launched on Linux operating system Ubuntu. We created a webserver called “IndiaBioDb” using HTML, PHP and CSS that support wide range of screens including smart phone, iPhone, iPad and desktops. This server have number of user-friendly web interfaces to extract information using wide range of options; following is brief description of major interfaces.

#### A. SEARCH

This module allows the user to access or retrieve data effortlessly using the simple search option. User can provide query against any field of database such as method name, journal, year, funding agency, author, city, class, and webserver type. It also allow user to select fields to be displayed, such as email of the author, PMID, source, description, category, link. Once the user types the keyword and selects the required fields to be displayed, the output page will provide all the relevant information associated with the keyword. User can click on the hyperlink of the software to access it.

#### B. BROWSE

We integrate browsing interface in web server that allows a user to extract information based on different fields like Class, Corresponding author, Institute/City, Journal, and Year. Based on the data compiled, we have provided a few entries in each section. User can click on the numbers written in front of the name that will allow uses to access all the related information.

#### C. Information

In this option, we have acknowledged some of the highly cited Indian bioinformaticians and also provided statistics page, where all the statistics of the data present in the database are provided.

Figure 1 represents the home page of IndiaBioDb, which is freely available and can be viewed on all browsers. It can be accessed at https://webs.iiitd.edu.in/raghava/indiabiodb/index.html.

**Figure 1.**
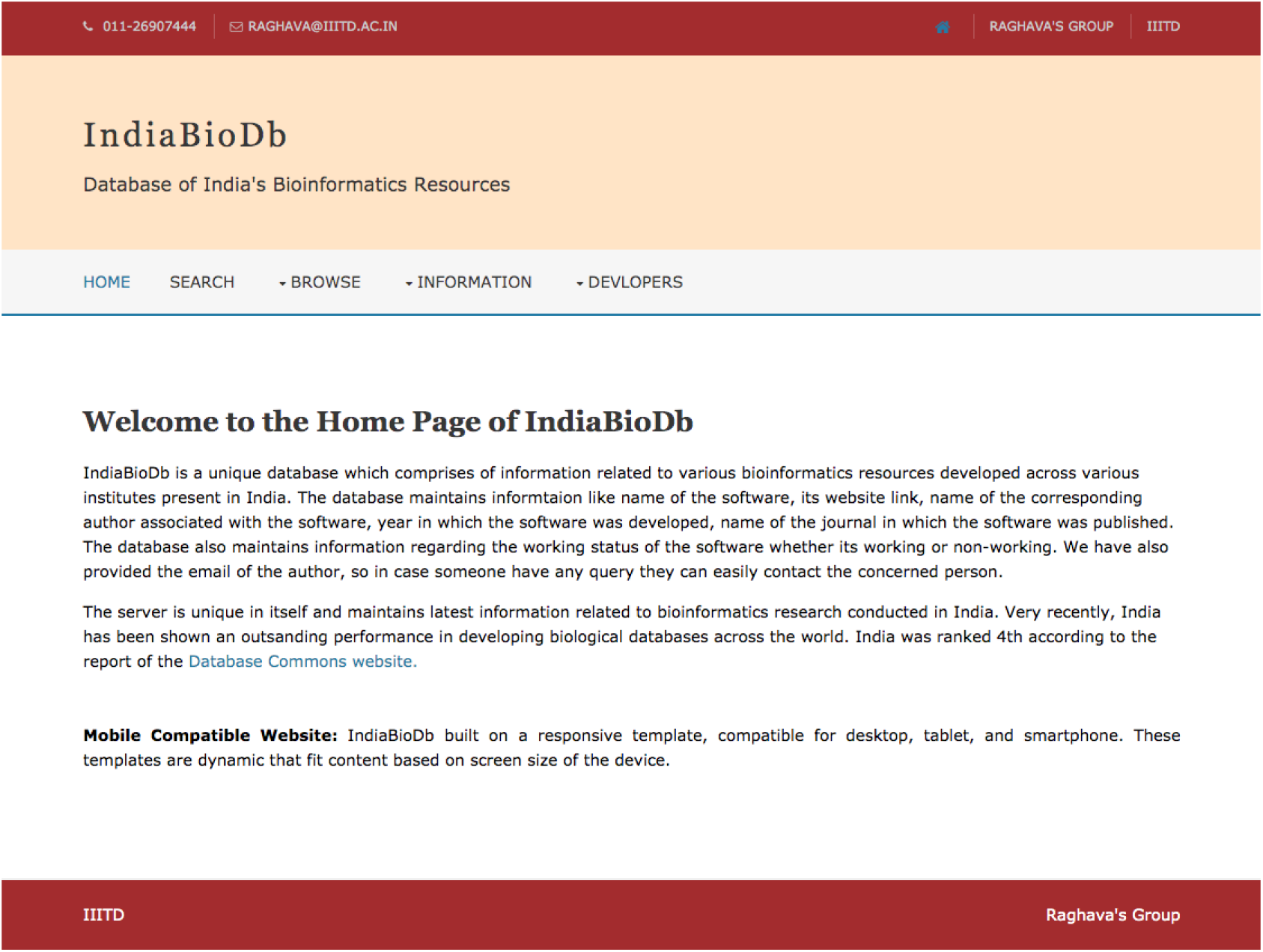
Screenshot of IndiaBioDb home page.

### Mobile App

We have also developed a Mobile App using Java to retrieve or extract information from our database. The app is very user-friendly and interactive; user can access all the bioinformatics data related to India from anywhere even without Internat. The ‘.apk’ file is provided at the database website for download as well as on Google Play Store.

## Results

Last two decades has seen a tremendous growth in the field of bioinformatics and computational biology. India is one of the countries who has contributed immensely in this field. Large number of databases, prediction algorithms and software has been developed and maintained by the Indian scientists. These web servers has been supported by various government and private bodies. BTISNet is one of the oldest and prominent bodies who is working in the growth of bioinformatics research in India. Table 1 provides statistics of bioinformatics resources developed at some of the centers of BTISNet. Individual principle investigators from various other instates institutes also developed number of resources. Major institutes outside BTISNet includes National Centre for Biological Sciences, Institute of Bioinformatics and Institute of Genomics and Integrative Biology.

**Table 1:** Shown major institute/centre supported by DBT under BTISNet and number of functional resources developed by these centres.

| Sr. No. | Centres | Number |
| --- | --- | --- |
| 1 | Institute of Microbial Technology | 165 |
| 2 | Indian Institute of Sciences | 30 |
| 3 | Indian Institute of Technology, Delhi | 20 |
| 4 | NIPGR, New Delhi | 10 |
| 5 | University of Pune | 9 |

It has been shown in a recent study that more than a third (34%) of the most cited papers in science were bioinformatics papers [16]; bioinformatics papers got 31-fold more citations than non-bioinformatics papers. It indicates importance bioinformatics research or bioinformatics resources at national and international level. We have compiled highly cited bioinformatics resources developed by Indian community (Table 2). Recently, number papers has been published on bioinformatics resources developed in India that includes Som et al., [3], Deobagkar et al., [10], Yadav and Mohanty [7], and Kale et al., [9].

**Table 2:**
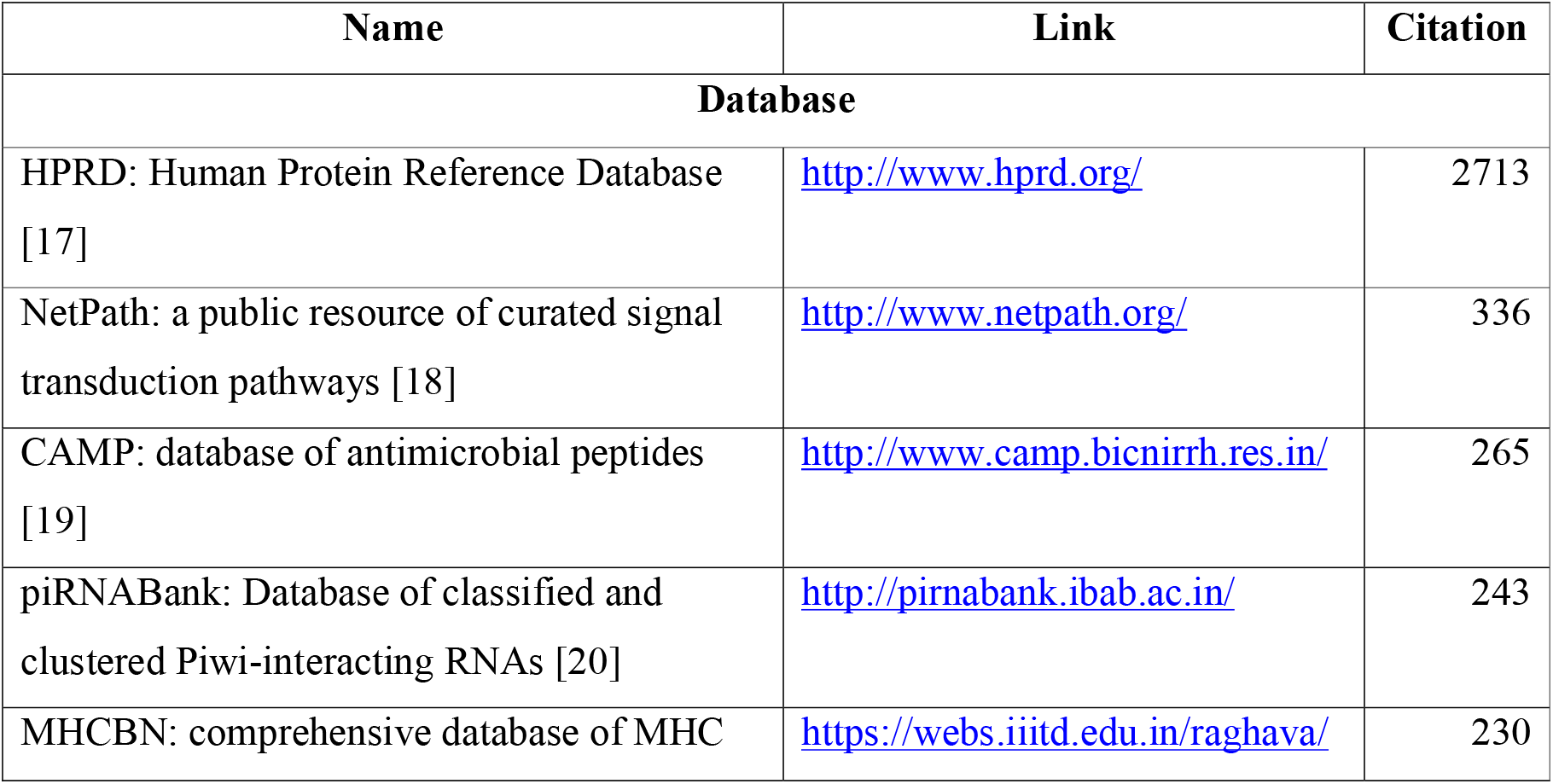
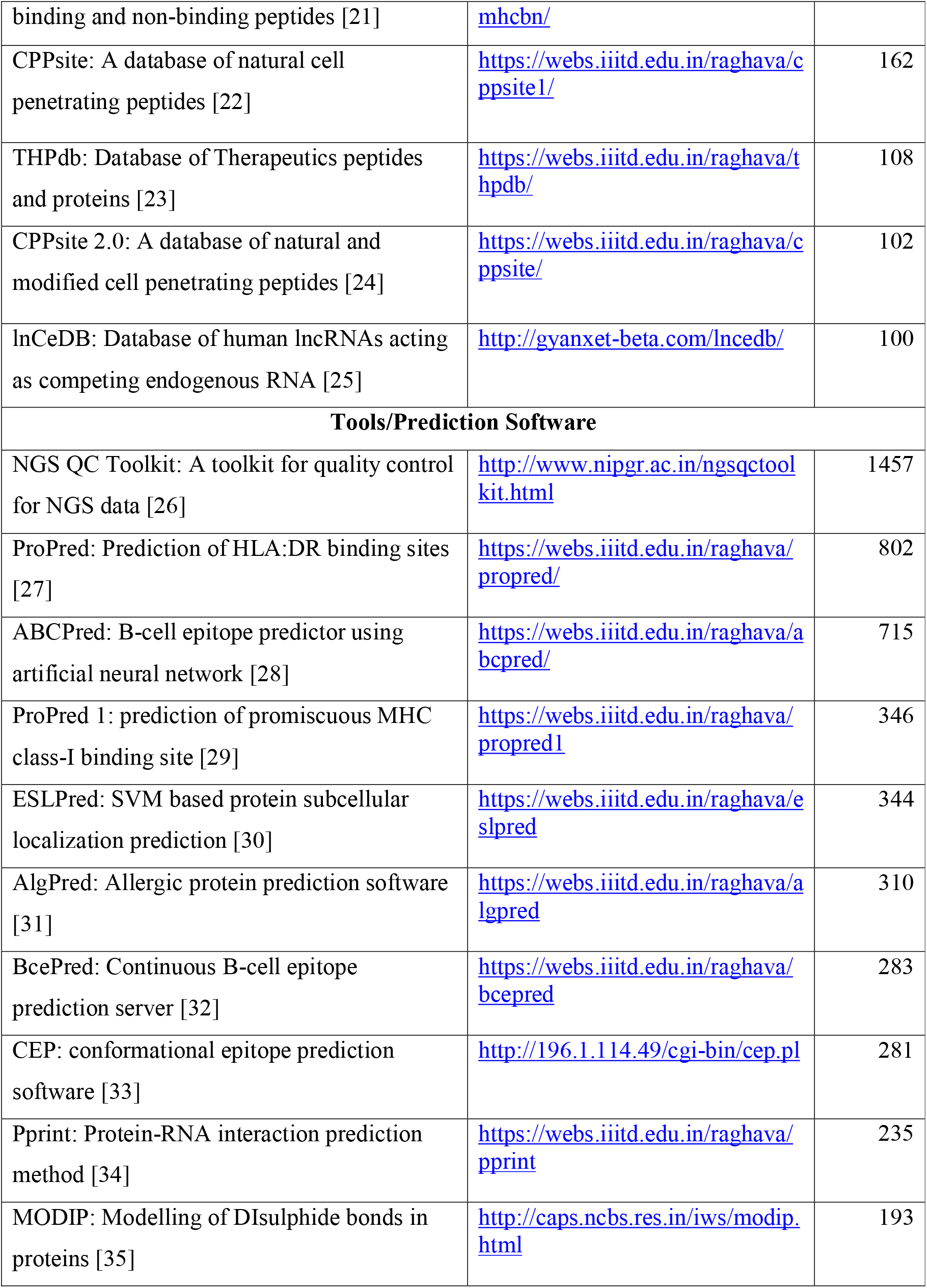
List of highly cited bioinformatics resources developed by Indian Bioinformaticians.

In order to highlight the contribution of Indian bioinformatics, we have compiled the data of functional resources, prediction methods and software developed and maintained in India. The data has been analysed by various means such as number of publications in the year, developing author, institute where the resource is developed and maintained, funding body supporting the institute and the resources. We grouped the articles published year-wise since 2001 and observed number of publications increased over the years (Figure 2). This show increasing interest of scientific community in bioinformatics in India. The analysis shows that Indian researchers developed several resources and most of these resources are functional. Statistics for the last five years is shown in Figure 2(A); complete statistics is provided in Supplementary Table S1. We grouped these resources in seven categories based on function; (i) Protein Structure; (ii) Protein Function; (iii) Vaccinomics; (iv) Genomics; (v) BioDrugs; (vi) Interactome and (vii) Cheminformatics. A maximum number 169 web servers were observed in the field of genomics. Protein Function and Protein Structure categories stand at second and third rank with 104 and 95 publications, respectively.

**Figure 2.**
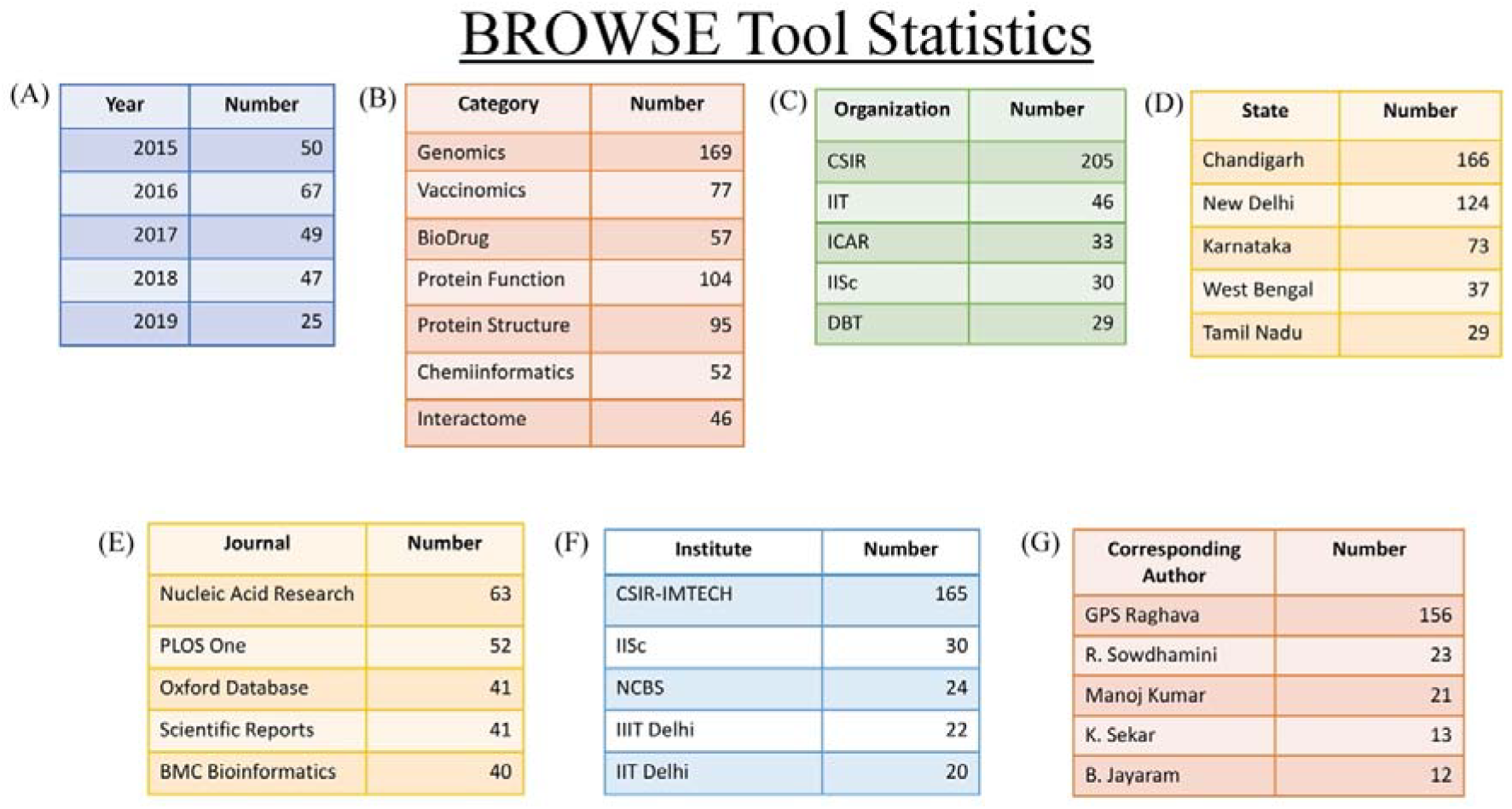
Statistics of various classes present in BROWSE tool.

Department of Biotechnology is one of the pioneers and oldest funding agencies promoting bioinformatics research in India since the early 1980s, which can be seen by the establishment of BTIS centres all across the country. Apart from BTIS centres, DBT has funded several individual projects as well as multi-institutional consortium projects such as the development of the NRRD database, a national database on *Mycobacterium tuberculosis*, database maintaining various information on Mango, etc. Apart from DBT, number of funding agencies (e.g., ICMR, ICAR, UGC) are also promoting bioinformatics research and funded numerous projects. It was observed CSIR institutes developed 205 web servers and IIT’s developed 46 resources, ranked first and second respectively. have been developed. Agricultural bioinformatics is one of the hot topics currently in India, and a number of bioinformatics research projects have been completed in this area. Some of them include the complete sequencing of the rice genome, tulsi genome, etc. ICAR is one of the bodies which is highly active in agricultural research, and currently, 33 web-based services are operational at various institutes affiliated to ICAR. Figure 2(C) provides statistics for the top five agencies; however, complete statistics is provided in Supplementary Table S2. We found that the maximum number of servers has been developed at Chandigarh with a total of 166, which is followed by 124 servers developed at Delhi. Karnataka stands at the third position with the total number of 73 servers. Figure 2(D) provides statistics for the first five states/area where scientists have developed a number of bioinformatics resources. Supplementary Table S3 provided detail figures for various states/areas.

Quality of research is paramount for any new scientific discovery, and those high quality research is regularly published in high-quality peer-reviewed journals. India is one of the few countries which over the years, has published many high-quality publications in peer-reviewed journals. 63 articles have been published in Nucleic Acid Research, 52 in PLOS One, 41 in Nature Scientific Reports, 40 in BMC Bioinformatics, and so on. These numbers show the quality of work done by Indian scientists and how their work has made an impact and contribution in the field of bioinformatics research. The number of publications is much higher in comparison to many other countries; however, countries like the USA, China, Japan, and the UK are publishing more than India, and one of the possible reasons could be the number of groups working in the field of bioinformatics in these countries. A list of the top 5 journals and the number of papers published in these journals is given in Figure 2(E). Complete statistics of articles published is given in Supplementary Table S4.

It is important to acknowledge the contribution of the Indian institutes in the area of bioinformatics research. These resources are well maintained and fully functional in certain institutes, which is important to utilize full potential of these services. CSIR-IMTECH is the leading institute with around 165 web servers developed by various scientific research groups. The servers developed at IMTECH provide a number of facilities in different field including drug and vaccine design. As per database commons, CSIR-IMTECH, Chandigarh is a leading institution; ranked at 4th number in the world (http://bigd.big.ac.cn/databasecommons/stat#). IIIT Delhi is one of the leading and newly emerging institutes in the field of bioinformatics as around 22 web services have been developed in the last five years. Indian bioinformatics is climbing the higher ladder in the field of bioinformatics day by day, and one of the reasons for this is the vision of scientists for benefitting society with their research. Due to the increasing popularity of bioinformatics nowadays, many researchers have taken bioinformatics as their career opportunity; however, few research groups in the field have contributed significantly in the last 20 years. Therefore, we also performed an analysis based on the corresponding authors and reported their contributions below. G.P.S. Raghava, Head of Department at Centre for Computational Biology, IIIT Delhi (earlier at CSIR-IMTECH) has developed maximum number of resources. His group developed more than 156 web servers collectively at CSIR-IMTECH, Chandigarh, and now at IIIT Delhi in the various field of bioinformatics. R. Sowdhamini from NCBS, Bangalore contributed 23 resources in the field of structural and functional proteomics. Some of the other top bioinformaticians from India include B. Jayaram from IIT Delhi, Manoj Kumar for CSIR-IMTECH, Michael M. Gromiha from IIT Madras, Shandar Ahmad from JNU, New Delhi, Dinesh Gupta from ICGEB, New Delhi. Indian bioinformaticians have published their research work in high-quality impact factors journals such as NAR, Oxford Bioinformatics, Genome Research, Nature Scientific Reports, BMC journals, Frontier journals. List of leading five bioinformaticians along with the number of functional web servers developed by them is given in Figure 2(G), and a complete list of other bioinformaticians is provided in Supplementary Table S6.

### Comparison with existing resources

We compared our repository or database with already existing repositories like database Commons, OMICS tools, Nucleic Acid Research (NAR), bio.tools at https://bio.tools supported by ELIXIR. Database Commons maintains information only about databases developed all over the world. It does not provide any information related to prediction tools, visualization tools, or any other such tools. Also, the user has to perform multiple clicks to extract all the information, which is a cumbersome task. Similarly, OMICS tools emphasize mostly on ‘omics’ data-based tools. It maintains information about the tools developed using multi-omics data (genomics, transcriptomics, proteomics, and metabolomics). bio.tools provides list of software and databases developed all over the world; however, it’s difficult to get information about resources specifically developed by Indian researchers and maintained in India. Similar problem is also faced by other repositories like NCBI at https://ncbi.nlm.nih.gov/guide/all and Bioinformatics Software and Tools at https://bioinformaticssoftwareandtools.co.in/index.php.

BTISNet site maintain link to databases and prediction tool developed by centres funded by DBT under BTISNet (http://btisnet.gov.in/btisdatabase.html). There are number of issues with this site, firstly it do not provide any information on about resources developed by other institutes of India. Secondly, it is not updated and do not provide comprehensive information these resources. In addition, number of articles and reviews on bioinformatics resources developed in India do not provide any site from where these resources can be retrieved [3,36–43]. In our study, we tried to address the shortcomings mentioned above by providing all the information related to Indian bioinformatics tools at a single platform. IndiaBioDb provides all the information about a tool in a single click. The information present in the databases is updated and about the web-based services which are functional and not obsolete. The mobile app of the database allows the user to access information globally more easily. These features provide an edge to this study in comparison to previous studies.

### Case Study

If any student wants to pursue a Ph.D. in India in the field of Genomics, he/she can easily search all the Principle Investigators (PIs) working in that particular field. User can type “Genomics” in the SEARCH Tab and select the class option, or click on the numbers in front of the class “Genomics” provided in the BROWSE tab. All the information related to the class “Genomics” such as names of tools, year and journal in which it is published, etc. will appear. In order to narrow down the list of PIs, user can further apply various filters such as the city or the institute where particular PI is working, quality of paper PI is publishing, etc.

## Discussion

In the past few decades, bioinformatics research has shown tremendous growth, and India is one of the leading countries which has taken an early step in establishing and promoting bioinformatics research. In this study, we have created a database IndiaBioDb, where we have compiled the information about the various databases and software developed in India. We have considered only those web servers which are functional as it will help wet lab scientists in performing various experiments, which will allow them to design their study more accurately with cheaper cost and within less time. User can also download the data from the databases and can develop novel or more sophisticated techniques which will help in better biological prediction. The database comprises of various information related to a web server such as the author who developed it, the institute where it has been maintained and its location (city and state). It also provides information related to the class of work it belongs to, the agency which has funded that institute, email of the corresponding authors, year in which the server was developed, journal in which the paper was published, link, and description of the software. The complete architecture of the database IndiaBioDb is shown in Figure 3.

**Figure 3.**
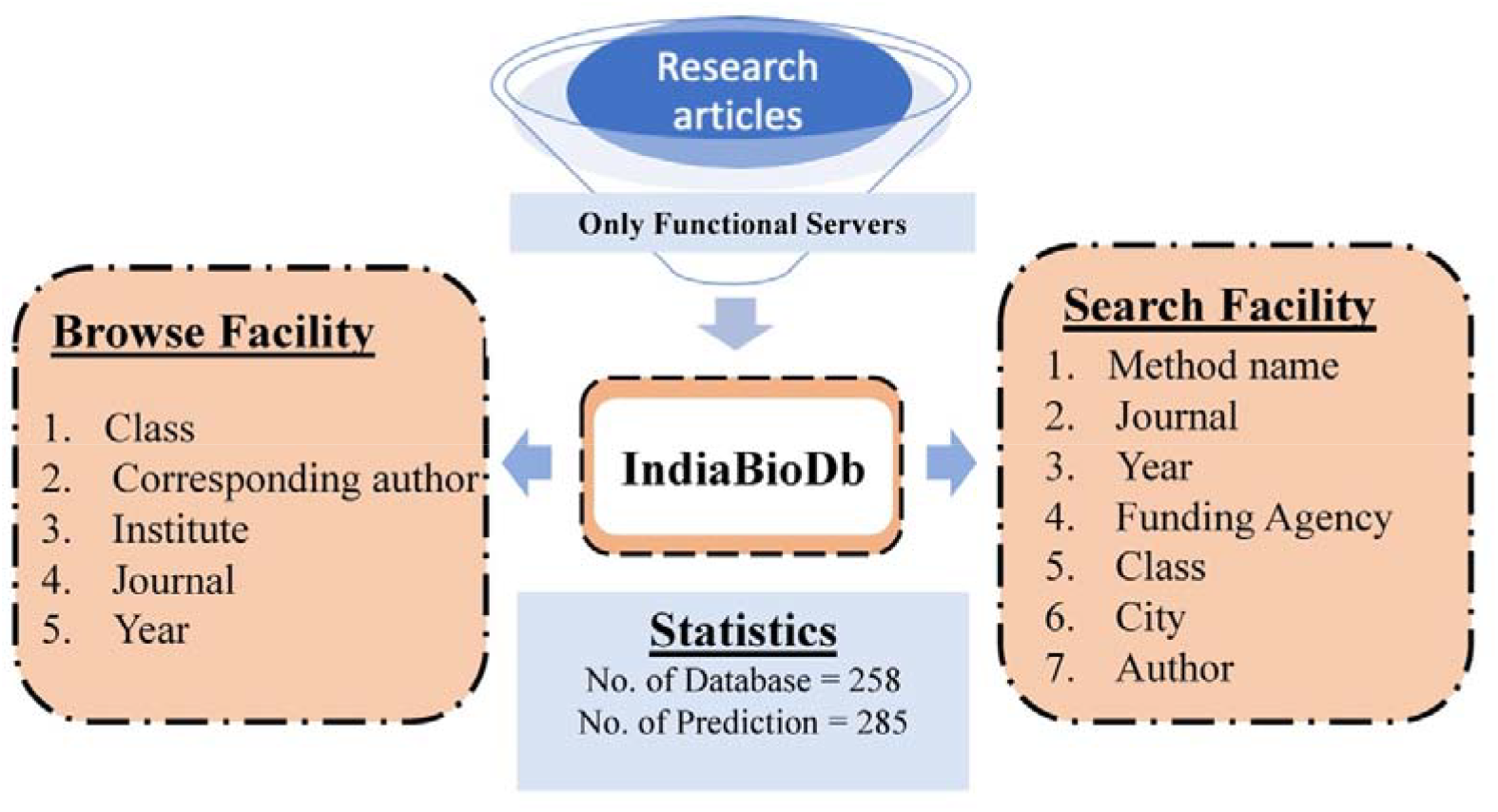
Architecture of IndiaBioDb.

There is a need to create more awareness among budding scientists as many of the states still doesn’t have bioinformatics course in their syllabus or research institutes for performing research. There is a requirement of more funds, infrastructure, job creation, and collaboration at a large scale to develop new databases and software which could help in solving real-life problems. In order to increase awareness among the students, we have created a repository where one can find all the details on the bioinformatics research done in India. The repository provides comprehensive information about different databases and software. User can visit our website or download the app for easy access to the data which is freely available. We believe that this work will allow scientists to use these web servers for their research purpose which will not only promote Indian bioinformatics also it will encourage the students to develop their career in this particular field.

## Conflict of Interest

The authors declare that they have no conflict of interest.

## Author’s Contribution

AA, AS, AD and PA collected, compiled and curated the data. AA, and PA performed the analysis and created the figures and tables. PA, and VK, developed the web interface. CM developed the mobile app. PA, AA, and GPSR wrote the manuscript. GPSR conceived the idea and supervised the project. All authors read and approved the final paper.

## Acknowledgement

Authors are thankful to J.C. Bose National Fellowship, Department of Science and Technology (DST), Government of India, Department of Science and Technology (DST-INSPIRE), University Grants Commission (UGC), Govt. of India and Indraprastha Institute of Information Technology, New Delhi, for fellowships and the financial support.

## Funding Information

This work was supported by J.C. Bose National Fellowship, Department of Science and Technology, Govt. of India.

