## Supplementary file for "A repository of web-based bioinformatics resources developed in India"

Prof. Gajendra P.S. Raghava

Head of Department of Computational Biology,

Indraprastha Institute of Information Technology

New Delhi 110020, India.

**Supplementary Data**

**Table S1: Year wise number of publications from India.**

| Year | Number |
| --- | --- |
| 2002 | 4 |
| 2001 | 3 |
| 2003 | 8 |
| 2004 | 18 |
| 2005 | 17 |
| 2006 | 14 |
| 2007 | 17 |
| 2008 | 22 |
| 2009 | 21 |
| 2010 | 25 |
| 2011 | 19 |
| 2012 | 40 |
| 2013 | 35 |
| 2014 | 62 |
| 2015 | 50 |
| 2016 | 67 |
| 2017 | 49 |
| 2018 | 47 |
| 2019 | 25 |

**Table S2: List of various Indian funding agencies and institutes maintaining different number of web servers.**

| Funding Agency | Number |
| --- | --- |
| CSIR | 205 |
| IIT | 46 |
| ICAR | 33 |
| IISc | 30 |
| DBT | 29 |
| IIIT | 26 |
| NCBS | 24 |
| JUIT | 11 |
| IISER | 10 |
| ICMR | 8 |
| University of Delhi | 7 |
| ISI | 6 |
| IBAB | 5 |
| IOB | 5 |
| IACS | 4 |
| ACTREC | 4 |
| VIT | 3 |
| CDAC | 2 |
| DTU | 2 |
| JNU | 2 |
| OTHERS | 82 |

**Table S3: List of various Indian states and territories maintaining different number of web servers.**

| State/Area | Number |
| --- | --- |
| Chandigarh | 166 |
| New Delhi | 124 |
| Karnataka | 73 |
| West Bengal | 37 |
| Tamil Nadu | 29 |
| Telangana | 25 |
| Maharashtra | 21 |
| Uttar Pradesh | 16 |
| Madhya Pradesh | 15 |
| Himachal Pradesh | 11 |
| Gujrat | 4 |
| Kerala | 4 |
| Odisha | 4 |
| Pondicherry | 3 |
| Punjab | 3 |
| Bihar | 2 |
| Assam | 2 |
| Uttarakhand | 2 |
| Andhra Pradesh | 1 |
| Assam Telangana | 1 |
| Jammu and Kashmir | 1 |

**Table S4: Number of research publications from India in international peer-reviewed journals.**

| Journals | Number |
| --- | --- |
| Nucleic Acids Research | 63 |
| PLOS One | 52 |
| Database: The Journal of Biological Databases and Curation | 41 |
| Scientific Reports | 41 |
| BMC Bioinformatics | 40 |
| Bioinformatics | 29 |
| Bioinformation | 24 |
| BMC Genomics | 11 |
| BMC Research Notes | 10 |
| Biology Direct | 9 |
| Genomics | 8 |
| Proteins | 8 |
| Genomics, Proteomics & Bioinformatics | 7 |
| In Silico biology | 7 |
| Protein and peptide letters | 7 |
| Frontiers in Microbiology | 6 |
| Methods in Molecular Biology | 6 |
| Bioinformatics and biology insights | 5 |
| Biodata Mining | 5 |
| Journal of Translational Medicine | 5 |
| Chemical biology and drug design | 4 |
| Frontiers in genetics | 4 |
| Journal of biosciences | 4 |
| Omics | 4 |
| The Journal of Biological Chemistry | 4 |
| 3 Biotech | 3 |
| Amino acids | 3 |
| Bioichimica et biophysics acta | 3 |
| Computers in biology and medicine | 3 |
| F1000 research | 3 |
| Frontiers in Immunology | 3 |
| Frontiers in plant science | 3 |
| Gene | 3 |
| Interdisciplinary Sciences: Computational Life Sciences | 3 |
| Journal of Computational Chemistry | 3 |
| Journal of molecular biology | 3 |
| Peer J | 3 |
| Systems and synthetic biology | 3 |
| Acta crystallographica section D biological crystallography | 2 |
| Biochemical and biophysical research communications | 2 |
| BMC Pharmacology | 2 |
| Computational Biology and Chemistry | 2 |
| FEBS letters | 2 |
| Frontiers in Pharmacology | 2 |
| Journal of bioinformatics and computational biology | 2 |
| Journal of biomedical informatics | 2 |
| Journal of biomolecular structure and dynamics | 2 |
| Journal of chemical information and modeling | 2 |
| Journal of Molecular Modeling | 2 |
| Journal of theoretical biology | 2 |
| Protein engineering,design and selection | 2 |
| Protein science | 2 |
| Others | 73 |

**Table S5:** **Number of research publications from various Indian institutions.**

| Institutes | Number |
| --- | --- |
| CSIR - Institute of Microbial Technology (CSIR-IMTECH) | 165 |
| Indian Institute of Science (IISc) | 30 |
| National Center for Biological Sciences (NCBS) | 24 |
| Indraprastha Institute of Information Technology (IIITD) | 22 |
| Indian Institute of Technology (IITD) | 20 |
| ICAR - Indian Agricultural Statistics Research Institute (IASRI) | 18 |
| CSIR - Institute of Genomics and Integrative Biology (CSIR-IGIB) | 12 |
| DBT - International Centre for Genetic Engineering and Biotechnology (ICGEB) | 11 |
| Indian Institute of Technology (IITM) | 11 |
| Jaypee University of Information Technology (JUIT) | 11 |
| DBT - National Institute of Plant Genome Research (NIPGR) | 10 |
| University of Pune | 9 |
| Indian Institute of Science Education and Research (IISERB) | 8 |
| CSIR - Centre for DNA Fingerprinting and Diagnostics (CSIR-CDFD) | 7 |
| University of Delhi (DU) South Campus | 7 |
| Institute of Bioinformatics (IOB) | 6 |
| Indian Statistical Institute (ISI) | 6 |
| Bose Institute | 5 |
| DBT - National Institute of Immunology (NII) | 5 |
| Institute of Bioinformatics and Applied Biotechnology (IBAB) | 5 |
| Advanced Centre for Treatment Research and Education in Cancer (ACTREC) Tata Memorial Centre | 4 |
| CSIR - Indian Institute of Chemical Biology (CSIR-IICB) | 4 |
| Indian Association for the Cultivation of Science (IACS) | 4 |
| Indian Institute of Technology (IITKGP) | 4 |
| University of Hyderabad | 4 |
| CSIR - Indian Institute of Chemical Technology (CSIR-IICT) | 4 |
| Others | 131 |

**Table S6:** **List of Indian bioinformaticians and the number of web servers they developed.**

| Corresponding Authors | Number |
| --- | --- |
| Gajendra PS Raghava | 156 |
| Ramanathan Sowdhamini | 23 |
| Manoj Kumar | 21 |
| K Sekar | 13 |
| B Jayaram | 12 |
| Dinesh Gupta | 11 |
| A R Rao | 9 |
| Michael Gromiha | 9 |
| Vineet K Sharma | 8 |
| Tiratha Raj Singh | 7 |
| Dinesh Kumar | 6 |
| Manish Kumar | 6 |
| Nagasuma Chandra | 6 |
| Debasisa Mohanty | 5 |
| Durai Sundar | 5 |
| Vinod Scaria | 5 |
| N Srinivasan | 4 |
| Akhilesh Pandey | 4 |
| Hampapathalu A Nagarajaram | 4 |
| Kshitish K Acharya | 4 |
| Srinivasan Ramachandran | 4 |
| Urmila Kulkarni Kale | 4 |
| Sudipto Saha | 4 |
| Amit Kumar | 3 |
| G N Sastry | 3 |
| Gitanjali Yadav | 3 |
| Jayashree Ramana | 3 |
| Jayprokas Chakrabarti | 3 |
| Mukesh Jain | 3 |
| Ramasubbu Sankararamakrishnan | 3 |
| Nita Parekh | 3 |
| Vaibhav Vindal | 3 |
| Others | 206 |
